# Shedding light on the embryo:endosperm balance diversity that pseudogamy can achieve in the polyploid apomictic Psidium cattleyanum (Myrtaceae, Myrteae)

**DOI:** 10.1101/2025.07.08.663781

**Authors:** Claudia Da Luz-Graña, Magdalena Vaio, Joerg Fuchs, Alejandra Borges, Gabriela Speroni

## Abstract

Apomixis is defined as the asexual mode of reproduction through seeds. Nearly all gametophytic apomictic species are polyploid. *Psidium cattleyanum*, a polyploid species with various described ploidy levels, exhibits apomixis and requires effective pollination for fruiting (pseudogamy). The aim of this study was to determine the frequency of apomixis and sexuality in *P. cattleyanum* among heptaploid and octoploid accessions, and to analyze the maternal and paternal contribution to endosperm formation. Flow cytometry was employed on mature seeds obtained from hand-pollinated crosses between two octoploid and two heptaploid accessions to determine the ploidy levels of embryo and endosperm. Within the 492 analyzed seeds the ploidy balance of embryo:endosperm ranged from the expected 2:5 (64.2%, pseudogamous apomixis) and 2:3 (4.7%, sexuality involving reduced gametophytes), to unexpected values, including 1:3 (0.4%),), 2:6 (2.4%), 2:7 (0.2%), 3:5 (8.8%), 2:3:5 (18.5%) and 2:4:6 (0.8%), associated with polyhaploid parthenogenesis, pseudogamous apomixis involving reduced and unreduced gametophytes, and female gametophytes with two or three polar nuclei in the central cell. The interpretations and implications of these results for apomictic polyploid species and their consequences for wild populations are discussed.

## INTRODUCTION

Polyploidy is defined as the presence of more than two sets of chromosomes in a nucleus and is recognized as one of the most important evolutionary forces in the phylogeny of angiosperms (Ramsey & Schemske, 1998; Soltis et al., 2015; Horandl, 2022; Heslop-Harrison et al., 2023). Genomic studies in the species *Amborella trichopoda* determined that a duplication event preceded the origin of all angiosperms (Amborella Genome Project, 2013). Therefore, all existing angiosperm species are paleopolyploids descending from a common ancestor that experienced at least one genome duplication (Jiao et al., 2011; Amborella Genome Project, 2013; Chanderbali *et al*., 2022). Polyploid formation can occur in various ways, most often through meiotic failures during sporogenesis that generate unreduced gametophytes. The male and/or female gametes of these gametophytes maintain the chromosome number of the species, giving rise to polyploid progenies after fertilization (Bretagnolle & Thompson, 1995; Ramsey & Schemske, 1998). Several authors have highlighted the close relationship between polyploidy and gametophytic apomictic reproductive systems (Hojsgaard & Hörandl, 2019; Hojsgaard & Pullaiah, 2022). Almost all apomictic species are polyploids (Nogler, 1984; Fei *et al*., 2019) and apomixis occurs in at least 1469 species (Hojsgaard & Pullaiah, 2022) belonging to 300 genera of 78 families of angiosperms (Cornaro *et al*., 2023). Apomixis is a form of asexual reproduction through seed formation (Asker & Jerling, 1992) that successfully overcomes the reproductive barriers between the parental genomes of newly formed polyploids and the meiotic irregularities resulting from the increase in genome numbers (Fei *et al*., 2019; Barke *et al*., 2020). It has been reported that in populations of species with different ploidy levels, odd cytotypes encounter difficulties in sexual reproduction and are more strongly associated with asexual reproduction (Kolář *et al*., 2017). Apomixis can be divided into two categories, gametophytic or sporophytic, depending on the presence or absence of embryo sac formation, respectively. Gametophytic apomixis can be further subdivided into apospory and diplospory, based on the origin of the cells involved in the formation of the female gametophyte (Nogler, 1984; Asker & Jerling, 1992; Bicknell & Koltunow, 2004). In this type of apomixis, female gametophytes with unreduced egg cells are regularly formed, where parthenogenetic development competes with fertilization (Dickinson *et al*., 2007), resulting in higher ploidy levels than their parental organisms. In approximately 90% of apomictic species, parthenogenesis excludes one of the processes of double fertilization, which is the fusion of a male gamete with the egg cell. However, the fertilization of the central cell is not necessarily nullified (Nogler, 1984; Hojsgaard & Hörandl, 2019; Terzaroli *et al*., 2023). This process, known as pseudogamy, not only involves the fertilization of the central cell for the formation of endosperm but also acts as a trigger for embryo development through parthenogenesis (Asker & Jerling, 1992).

In sexual species, there is a ratio between maternal and paternal contribution of 2m:1p for endosperm development, as described by Johnston *et al*. (1980). This 2m:1p ratio was identified as necessary for successful development of the nutritive tissue. In cases where this contribution is altered, embryo development is compromised (Johnston *et al*., 1980; Nogler, 1984). However, in apomictic species, the ratio of maternal and paternal contribution in the endosperm can vary. In *Paspalum notatum*, an aposporic pseudogamous species, tetraploid cytotypes (4x) form endosperm with various ratios (4m:1p, 8m:1p) and produce seeds independent of the ploidy level of the pollinator (2x, 4x, 5x, 6x, and 8x) (Quarin, 1999). Similarly, in *Tripsacum dactyloides*, a diplosporic pseudogamous species with 4x cytotypes, seed formation was reported with pollen donors of 2x and 4x, resulting in endosperms with combinations of 8m:1p and 8m:2p (Grimanelli *et al*., 1997).

*Psidium* (Myrtaceae, Myrteae) is the fourth most species-rich genus of Myrtaceae and includes approximately 92 tree or shrub species, predominantly found in neotropical regions. *Psidium guajava* L. stands out within this genus due to its significant economic value as a fruit tree (Proença *et al*., 2022). Another notable fruit species within the genus is *Psidium cattleyanum* Sabine, distributed from northeastern Brazil to eastern Uruguay (Legrand, 1968; Legrand & Klein, 1977; Brussa & Grela, 2007; Speroni *et al*., 2012; Sobral *et al*., 2015). Due to its ornamental qualities and the delightful flavor of its fruits (Vignale & Bisio, 2005), *P. cattleyanum* Sabine has been introduced to various regions worldwide (Huenneke & Vitousek, 1990; Ellshoff *et al*., 1995; Koske & Gemma, 2006). In some of these regions, it has become a highly problematic invasive tree species (Huenneke & Vitousek, 1990; Koske & Gemma, 2006, Richardson & Rejmáenk, 2011). Two infraspecific forms are recognized, mainly differentiated by the color of their fruits: *P. cattleyanum* f. *cattleyanum,* which produces red fruits, and *P. cattleyanum* f. *lucidum,* that produces yellow fruits (Degener, 1939).

*Psidium cattleyanum* is a polyploid species with a basic chromosome number of x=11. No diploid materials have been identified, and the reported chromosome numbers range from triploid to decaploid and dodecaploid, and additionally the occurrence of some aneuploid materials was described (Atchinson, 1947; Singhal *et al*., 1985; Raseira & Raseira, 1996; Costa & Forni-Martins, 2006; Vázquez, 2014; Machado *et al*., 2020; Machado & Forni-Martins, 2022). Brazilian populations mostly consist of a single cytotype, although mix-ploidy populations with up to five different ploidy levels have also been reported (Machado *et al*., 2020; Machado & Forni-Martins, 2022). The origin of the different cytotypes in the species is unknown and both, allopolyploid and autopolyploid origin, have been suggested based mainly on cytogenetic studies (Costa, 2009; Vázquez, 2014; Machado & Forni-Martins, 2022). *P. cattleyanum* is easily propagated by seeds, has a high germination rate (Franzon *et al*., 2009) and the progenies present a high morphological uniformity among themselves and with respect to the mother plant (Raseira & Raseira, 1996; Vignale & Bisio 2005). Apomixis was observed in other species of the Myrtaceae family, where evidence of apospory and adventitious embryony was found (Johnson, 1936; Gurgel & Soubihe Sobrinho, 1951; Nic Lughadha & Proença, 1996; Thurlby *et al*., 2012). *P. cattleyanum* is the first species of Myrtaceae where diplosporic apomixis was reported. Ontogenetic examinations of the ovule, conducted on heptaploid and octoploid materials, revealed a very low frequency of meiosis of the mother cell of the megaspore and the absence of aposporic embryo sacs and adventitious embryos (Souza-Pérez & Speroni, 2017). In the same study, it was additionally noted that fruiting occurs only with the arrival of viable pollen on the stigma, regardless of its origin, suggesting the presence of pseudogamy, as in most apomictic species. An interesting observation made by the authors of this study is the presence of three polar nuclei in several embryonic sacs (Souza-Pérez & Speroni, 2017). This observation could potentially impact the ploidy level of the central cell and, consequently, influence the balance of the embryo:endosperm ploidy levels.

In addition to the apomictic system, certain evidence of sexuality has also been found in *P. cattleyanum*. A study using SSR markers on three wild populations from Brazil revealed varying degrees of genetic variability in populations associated with different ploidy levels. The authors interpreted these findings as indicative of the coexistence of apomixis and sexuality (Machado *et al*., 2022). It is known that in facultative apomictic species a certain degree of sexuality remains (Bicknell & Koltunow, 2004; Reutemann *et al*. 2022; Hörandl, 2022), and both pathways can even coexist on the same plant (Nogler, 1984).

Flow cytometric seed screen (FCSS) is based on the determination of the different DNA contents / ploidy levels present in mature seeds and the balance that exists between them indicates the reproductive origin of the seeds (Matzk *et al*., 2000). FCSS discriminates between sexual and apomictic embryos, but also different mechanisms of apomictic reproduction and gamete characteristics (Krahulcová & Rotreková, 2010). A seed whose embryo originated sexually has a ploidy balance between embryo and endosperm of 2:3, due to the 2C (1C+1C) composition of the embryo and 3C (1C+1C+1C) of the endosperm. In the case of apomictic seeds, the embryo remains 2C (2C+0) but the central cell forming the endosperm is 4C (2C+2C+0) due to the lack of meiotic reduction and fertilization. Therefore, the embryo:endosperm balance is 2:4. If fertilization of the middle cell is necessary for endosperm formation (pseudogamy), it will be 5C (2C+2C+1C) and the embryo:endosperm balance will be 2:5 (Matzk *et al*., 2000). FCSS has been used successfully in several species such as *Hypericum perforatum* (Matzk *et al*., 2001) and *Cotoneaster integerrimus* (Macková *et al*., 2020); and genera such as *Crataegus*, *Mespilus* (Talent & Dickinson, 2007) and *Boechera* (Aliyu *et al*., 2010), allowing the pathway of embryo and endosperm formation to be determined (Loureiro *et al*., 2023).

Considering the polyploid and apomictic status of *Psidium cattleyanum*, the aims of this work were 1) Determining the frequency at which apomictic and sexual pathways occur in the seed formation in *P. cattleyanum*, 2) Comparing potential differences in the reproductive pathways between plants with even (8x) and odd ploidy (7x), and 3) Analyzing the maternal-paternal contributions to the formation of the endosperm.

For this, we used the FCSS on mature seeds obtained in different crosses between octoploid (2n=8x=88) accessions of the *lucidum* form (yellow fruits) and heptaploid (2n=7x=77) accessions of the *cattleyanum* form (red fruits). The interpretation of its reproductive origin was deduced from the balances in DNA content / ploidy level between the embryo and endosperm.

## MATERIALS AND METHODS

### Plant material

Seeds of *Psidium cattleyanum* were obtained through hand-pollinations between two octoploid accessions (2n=8x=88) of yellow-fruited plants (III5 and IV6) and two heptaploid accessions (2n=7x=77) of red-fruited plants (IV1 and IV7). The ploidy levels and chromosome numbers of these accessions were previously determined by Vázquez (2014). All plants are part of the ‘Program of Wild Fruits with Economic Potential’ (Facultad de Agronomía-Instituto Nacional de Investigación Agropecuaria) and are cultivated in the Experimental Field Station of Facultad de Agronomía (EEFAS, 31°19’S, 57°41’W), Dpto. Salto, Uruguay. These plants were selected based on the flavor of their fruits and early fruiting. Vouchers for *P. cattleyanum* f. *cattleyanum* IV1 (Speroni 1042) and IV7 (Speroni 1041), and *P. cattleyanum* f. *lucidum* III5 (Speroni 1037) and IV6 (Speroni 1040) were deposited in the Bernardo Rosengurtt Herbarium (MVFA), Facultad de Agronomía, Universidad de la República, Uruguay. Before hand pollination, pollen grain viability of each plant was assessed using a colorimetric test with 2,3,5 triphenyl tetrazolium chloride (TTC) (Shivanna & Rangaswamy, 1992). Based on the results obtained (32.81% in III5, 39.04% in IV6, 11.02% in IV1 and 1.15% in IV7), plant IV7 was excluded as a pollen donor due to its low pollen grain viability values.

### Hand pollinations treatments

Hand pollinations were conducted between individuals of the four accessions (III5, IV1, IV6 and IV7) of *P. cattleyanum.* Due to the lack of exact synchronization between the flowering periods of the plants, only a limited timeframe was available for the pollen transfer. Within this period, 3 to 17 randomly chosen flowers from each tree underwent emasculation one hour before anthesis (when the anthers are still indehiscent). Additionally, the stigma was isolated to prevent the introduction of foreign pollen grains. Four hours later, hand pollination was performed using pollen from another plant of the same ploidy (homoploidy pollination) or different ploidy (heteroploidy pollination) (Table 1). For self-pollination treatments, pollen from the flower itself was used without emasculating and bagged prior to anthesis to prevent contamination (Table 1). Fruits were monitored until maturity for seed extraction. The number of seeds obtained from each pollination treatment ranged between 59 and 373 per fruit, of which approximately 40 were used for cytometry analyses.

**Table 1.** Hand pollination treatments in heptaploid (7x) accessions (IV1 and IV7), and octoploid (8x) accessions (III5 and IV6) of *Psidium cattleyanum*.

| Pollen receptor | Pollen donor |  |  |
| --- | --- | --- | --- |
|  | III 5 (8x) | IV 6 (8x) | IV 1 (7x) |
| III 5 (8x) | self-pollination | homoploidy | heteroploidy |
| IV 6 (8x) | homoploidy | self-pollination | heteroploidy |
| IV 1 (7x) | heteroploidy | heteroploidy | self-pollination |
| IV 7 (7x) | heteroploidy | heteroploidy | homoploidy |

### Flow cytometric seed screen (FCSS)

The seed development pathway in *Psidium cattleyanum* was determined using the flow cytometric seed screen (FCSS) method as proposed by Matzk *et al*. (2000) with some modifications. A key difference is the use of single seed analysis instead of bulk seed analysis. Prior to the nuclei isolation, the seed coat was removed mechanically. The nuclei were isolated using the procedure and buffer described by Galbraith *et al*. (1983). Afterwards, the isolated nuclei were filtered through a 30 µm mesh and stained with propidium iodide (50 µg/mL final concentration). A Partec CyFlow® Space flow cytometer (Sysmex-Partec GmbH, Münster, Germany) was utilized, and the different cell populations were analyzed using the FloMax® program (Version 2.4d; Sysmex-Partec GmbH, Münster, Germany). This approach enabled the visualization of the relative DNA content for the nuclei of different seed tissues as distinct peaks in a histogram for each seed. The balance between peaks corresponding to the embryo and the endosperm allowed the interpretation of the seed development pathway. To precisely determine the relationship between the different peaks the position of the embryo peak (2C value) was manually set to channel 50 by adjusting the gain of the FL1 parameter. Additionally, mean values of each peak were manually determined after each measurement. Nuclei populations corresponding to the embryo were validated by determining the DNA content of leaf nuclei of plants with known ploidy, using *Lycopersicon esculentum* Mill. convar. infiniens Lehm. var. *flammatum* Lehm. ‘Stupicke Rane’ (2C=1.96 pg, IPK genebank accession number: LYC 418) as an internal standard. As an example, for seeds in which the embryo peak was placed to channel 50 and the corresponding endosperm peak was detected at FL1 channel 75, a 2:3 balance was considered. This balance represents a fertilization of the egg cell by a male gamete (1C+1C) and the resultant endosperm, corresponding to the fertilization of two polar nuclei by a male gamete (1C+1C+1C). Seeds where the endosperm peak was located at channel 125 were considered to have a balance of 2:5 (2C+0:2C+2C+1C). In both balances, 2:3 and 2:5, ploidy of the embryo and endosperm was determined by the presence of reduced and unreduced male and female gametes. Consequently, other balances were observed indicating different levels of ploidy in the embryo and endosperm, based on the reduced condition or not of the embryo sac and the male gametes participating in fertilization (Matzk *et al*., 2000). Results where the peaks were not easily assignable or the endosperm was not visualized were excluded from the analyses. To determine variations in ploidy level of the embryo we used the instrument settings automatically recorded by the FloMax® program. Changes in FL1 gain settings between runs were used to determine changes in ploidy levels.

### Data analysis

The proportion of apomictic and sexual seeds obtained from the different crosses was determined using a generalized linear model (GLM), assuming a binomial distribution. An F-test was performed to test fixed effects. Given the hierarchical structure of the cross design (ploidy, mother, pollen donor), orthogonal contrasts were performed to compare the means per group of plants: 1. 7x mothers vs 8x mothers, 2. within 7x mothers (IV1 vs IV7), 3. within 8x mothers (III5 vs IV6), 4. within 7x_IV1 mother (IV1×IV1 vs IV1 with others pollen donor), 5. within 7x_IV7 mother (IV7×IV1 vs IV1 with others pollen donor), 6. within 8x_III5 mother (III5×III5 vs III5 with others pollen donor) and 7. within 8x_IV6 mother (IV6×IV6 vs IV6 with others pollen donor). All these analyses were done using PROC GLIMMIX from SAS software (SAS, 2016, software for Windows version 9.4, SAS Institute Inc, Cary, NC).

## RESULTS

### Embryo:endosperm DNA content balances

The flow cytometry seed screen revealed a greater diversity in the pathways of embryo and endosperm formation in *Psidium cattleyanum* than expected. A total of 492 seeds were analyzed, of which 316 (64.2%) showed a 2:5 balance corresponding to an apomictic pathway with fertilization of the central cell of an unreduced embryo sac (pseudogamy), and involving a reduced male gamete. Only 23 seeds (4.7%) exhibited a 2:3 balance corresponding to events of sexuality between reduced gametophytes. The remaining seeds represented other DNA content balance of 1:3, 2:6, 2:7, and 3:5 (Table 2), corresponding to events of haploid parthenogenesis (1:3), diploid parthenogenesis (2:6 and 2:7), and sexuality (3:5). These events resulted from combinations involving a reduced or unreduced embryo sac fertilized by one or two reduced or unreduced male gametes to form the embryo and endosperm of each seed (Figure 1). In addition to these results, 95 seeds (19.3%) exhibited the presence of a third peak in the histogram, showing 2:3:5 or 2:4:6 balances. This is associated with unreduced apomictic embryo sacs with three polar nuclei in the central cell and the formation and coexistence of nuclei with two ploidy levels in the endosperm. Representative histograms of the main observed balances are shown in Figure 2.

**Figure 1.**
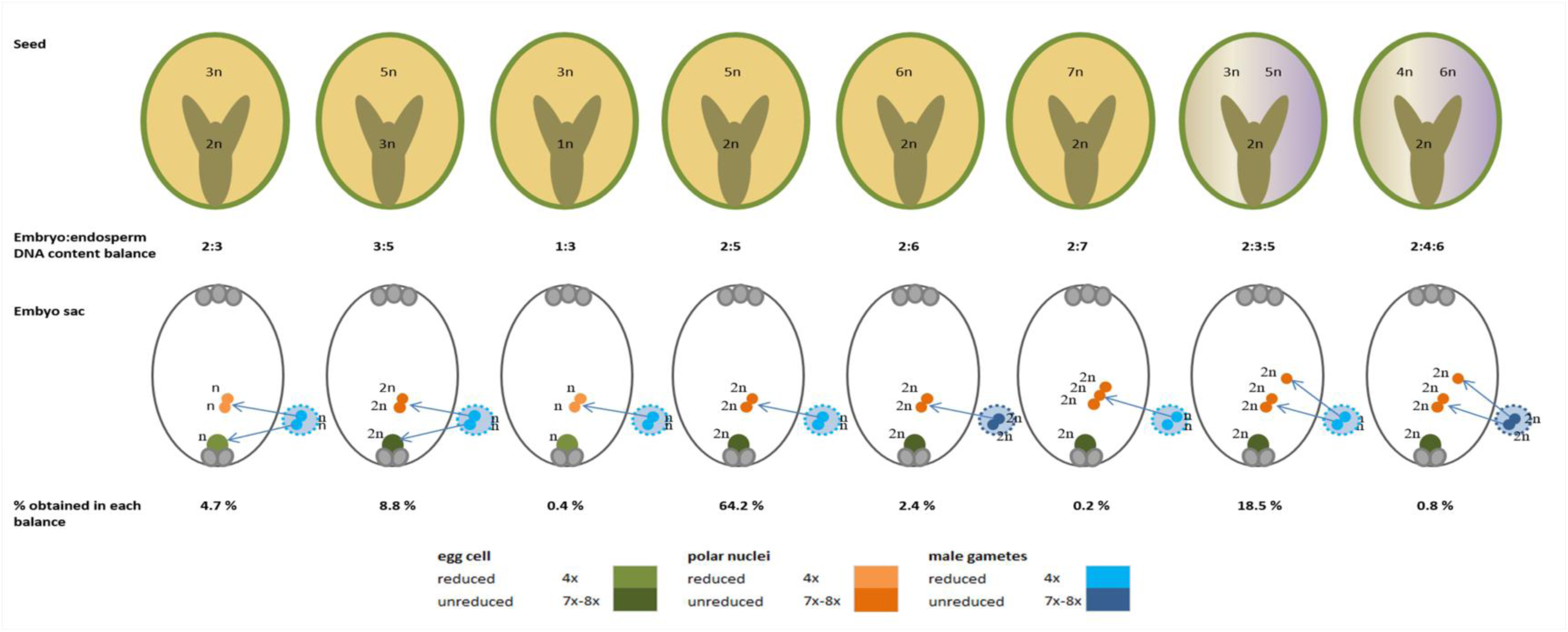
Scheme illustrating the different pathways of seed formation resulting in various embryo:endosperm balances (or embryo:endosperm:endosperm in the case of the presence of three polar nuclei) in *Psidium cattleyanum* seeds obtained from manual crosses of heptaploid and octoploid accessions analyzed by the flow cytometric seed screen (FCSS) method. The depicted combinations showcase the female gametophyte and male gametes contributing to the seed formations.

**Figure 2.**
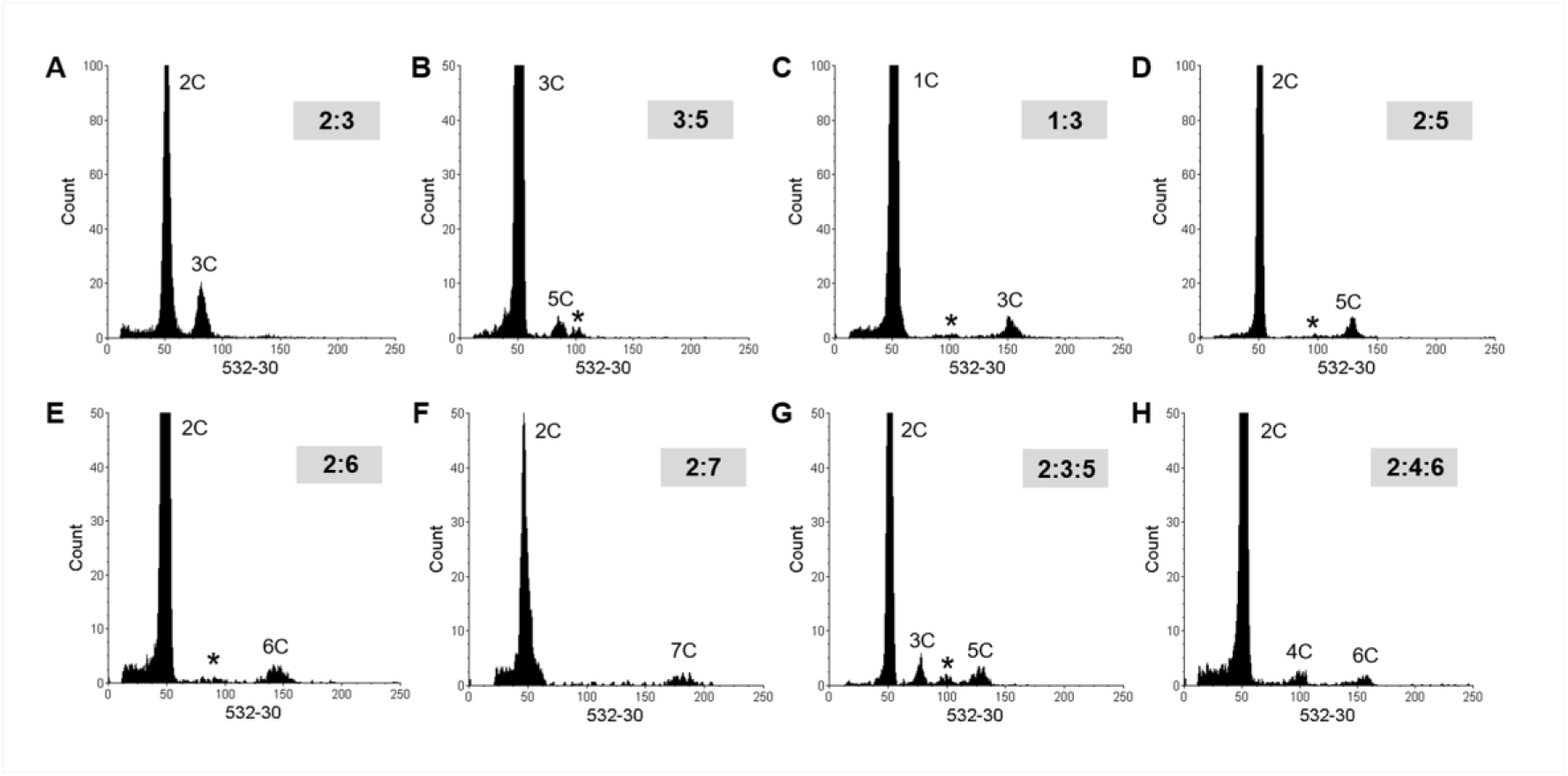
Representative histograms of flow cytometric seed screen (FCSS) measurements in *Psidium cattleyanum* showing the different DNA content balances between embryo : endosperm (or embryo : endosperm : endosperm) observed. Peaks correspond to embryo or endosperm nuclei in the G0/G1 cell cycle phase and to the balances: A 2:3 (sexual), B 3:5 (sexual), C 1:3 (apomictic), D 2:5 (apomictic), E 2:6 (apomictic), F 2:7 (apomictic), G 2:3:5 (apomictic) and H 2:4:6 (apomictic). The asterisks (*) in B, C, D, E, and G indicate embryo nuclei in G2 cell cycle phase. The corresponding fcs files were edited using the software FCS Express 7 Flow (version: 7.12.0007; De Novo Software). Note that in B and C the gain of the fluorescence parameter was changed to set the embryo peak to channel 50, see Materials & Methods.

**Table 2.**
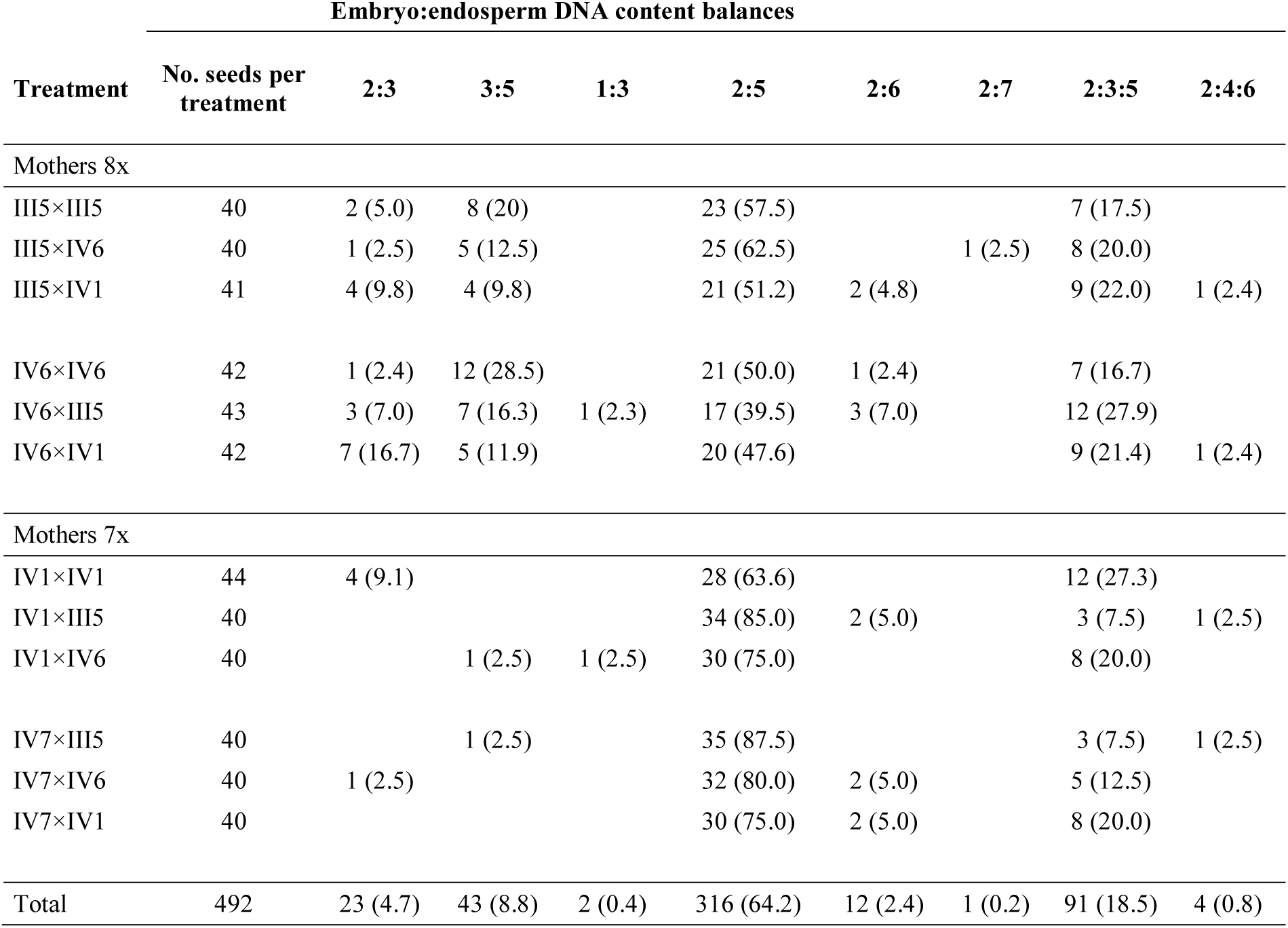
Number of seeds (percentage) of *Psidium cattleyanum* recorded in different embryo:endosperm DNA content balances for each hand pollination treatment. The relationship between embryo:endosperm (or embryo:endosperm:endosperm for cases where nuclei with two different ploidy levels coexist in the endosperm), the ploidy level (7x or 8x) of the plants used (III5: 8x, IV6: 8x, IV1: 7x, IV7: 7x) and the total number of seeds per treatment are shown.

| Embryo:endosperm DNA content balances |  |  |  |  |  |  |  |  |  |
| --- | --- | --- | --- | --- | --- | --- | --- | --- | --- |
| Treatment | No. seeds per treatment | 2:3 | 3:5 | 1:3 | 2:5 | 2:6 | 2:7 | 2:3:5 | 2:4:6 |
| Mothers 8x |  |  |  |  |  |  |  |  |  |
| III5×III5 | 40 | 2 (5.0) | 8 (20) |  | 23 (57.5) |  |  | 7 (17.5) |  |
| III5×IV6 | 40 | 1 (2.5) | 5 (12.5) |  | 25 (62.5) |  | 1 (2.5) | 8 (20.0) |  |
| III5×IV1 | 41 | 4 (9.8) | 4 (9.8) |  | 21 (51.2) | 2 (4.8) |  | 9 (22.0) | 1 (2.4) |
| IV6×IV6 | 42 | 1 (2.4) | 12 (28.5) |  | 21 (50.0) | 1 (2.4) |  | 7 (16.7) |  |
| IV6×III5 | 43 | 3 (7.0) | 7 (16.3) | 1 (2.3) | 17 (39.5) | 3 (7.0) |  | 12 (27.9) |  |
| IV6×IV1 | 42 | 7 (16.7) | 5 (11.9) |  | 20 (47.6) |  |  | 9 (21.4) | 1 (2.4) |
| Mothers 7x |  |  |  |  |  |  |  |  |  |
| IV1×IV1 | 44 | 4 (9.1) |  |  | 28 (63.6) |  |  | 12 (27.3) |  |
| IV1×III5 | 40 |  |  |  | 34 (85.0) | 2 (5.0) |  | 3 (7.5) | 1 (2.5) |
| IV1×IV6 | 40 |  | 1 (2.5) | 1 (2.5) | 30 (75.0) |  |  | 8 (20.0) |  |
| IV7×III5 | 40 |  | 1 (2.5) |  | 35 (87.5) |  |  | 3 (7.5) | 1 (2.5) |
| IV7×IV6 | 40 | 1 (2.5) |  |  | 32 (80.0) | 2 (5.0) |  | 5 (12.5) |  |
| IV7×IV1 | 40 |  |  |  | 30 (75.0) | 2 (5.0) |  | 8 (20.0) |  |
| Total | 492 | 23 (4.7) | 43 (8.8) | 2 (0.4) | 316 (64.2) | 12 (2.4) | 1 (0.2) | 91 (18.5) | 4 (0.8) |

### Apomixis vs. Sexuality

For statistical analysis of the balance between apomixis and sexuality in the different crosses, the observed DNA content balances were grouped based on their sexual or apomictic pseudogamous origin (with two or three polar nuclei). The frequency distribution of each origin showed a wide range starting from the absence of sexuality in crosses of IV1×III5 (7x×8x) and IV7×IV1 (7x×7x) to a maximum of 31% sexuality in the cross IV6×IV6 (8x×8x) (Figure 3). Among the seven orthogonal contrasts performed, only the one comparing the proportion of sexual seeds between ploidies (odd vs even ploidy) was statistically significant (p=0.0158). It was observed that out of 248 seeds analyzed from 8x mother plants, 24% were formed sexually, whereas for the 244 seeds from the odd 7x mother plants, only 3% originated through sexual reproduction. In the comparison of groups based on pollen donor plants (III5, IV6, and IV1), it was observed that the frequency distribution of seeds of sexual origin in each group is in all cases slightly above 10%, with no significant differences between them (p=0.05).

**Figure 3.**
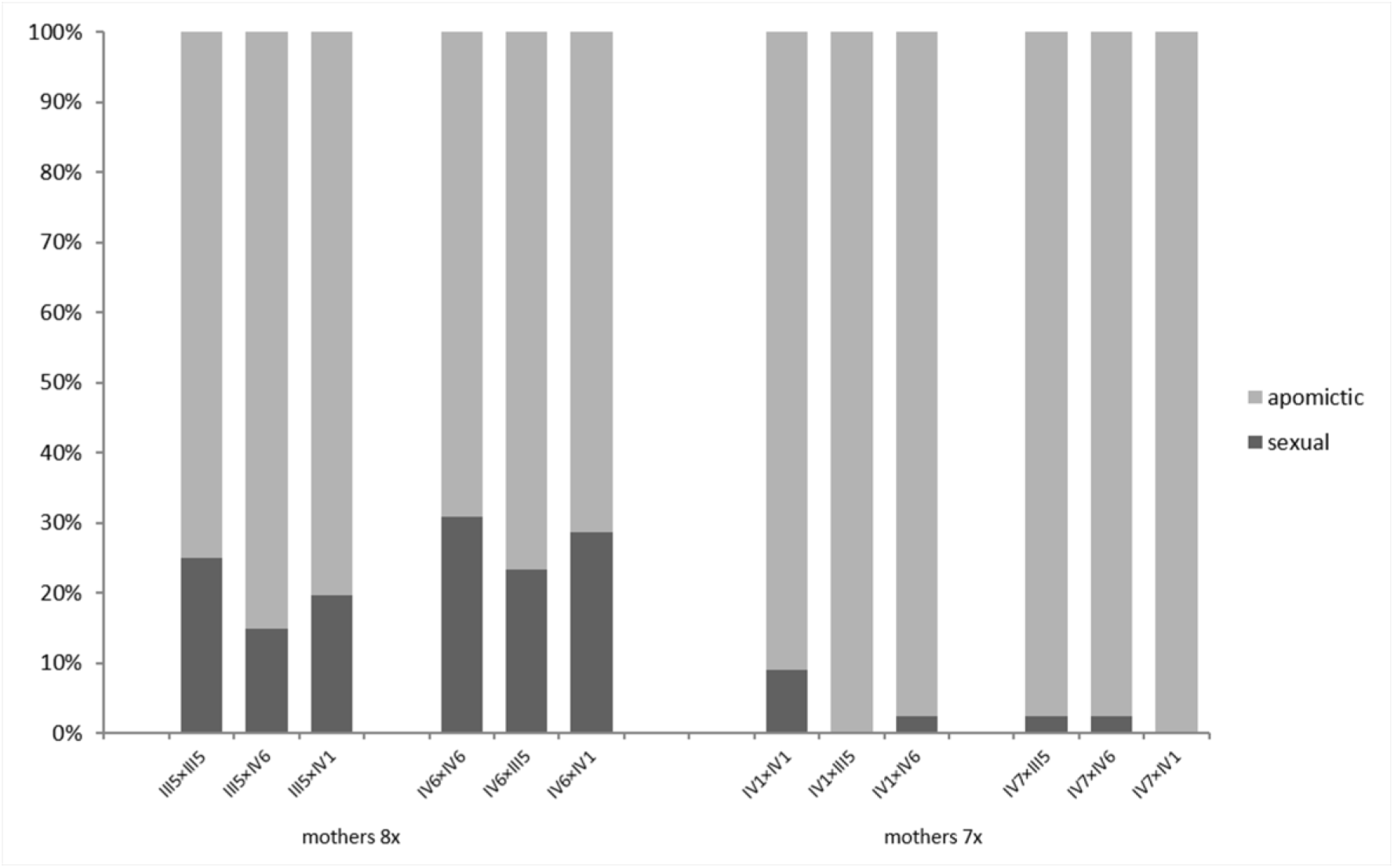
Frequency distribution of seeds of sexual or apomictic origin resulting from hand-pollinated crosses in *Psidium cattleyanum* considering the ploidy level (8x or 7x) of the mother plants. III5 and IV6 octoploid plants (yellow fruits); IV1 and IV7 heptaploid plants (red fruits).

### Ploidy Levels of Gametophytes, Embryo and Endosperm

The results obtained from DNA content analysis of the embryo and endosperm revealed that the majority of mother plants (94.9%) produced unreduced embryo sacs, while the majority of the involved pollen grains (96.8%) are reduced, including those generated by the 7x plant, which have a relative DNA content equivalent to 4x (Figure 4). The ploidy level of the embryo in most cases (89.8%) was the same as that of the mother (7x or 8x). In the remaining cases DNA contents for the embryo correspond to 4x, 11x, and 12x (Figure 4).

**Figure 4.**
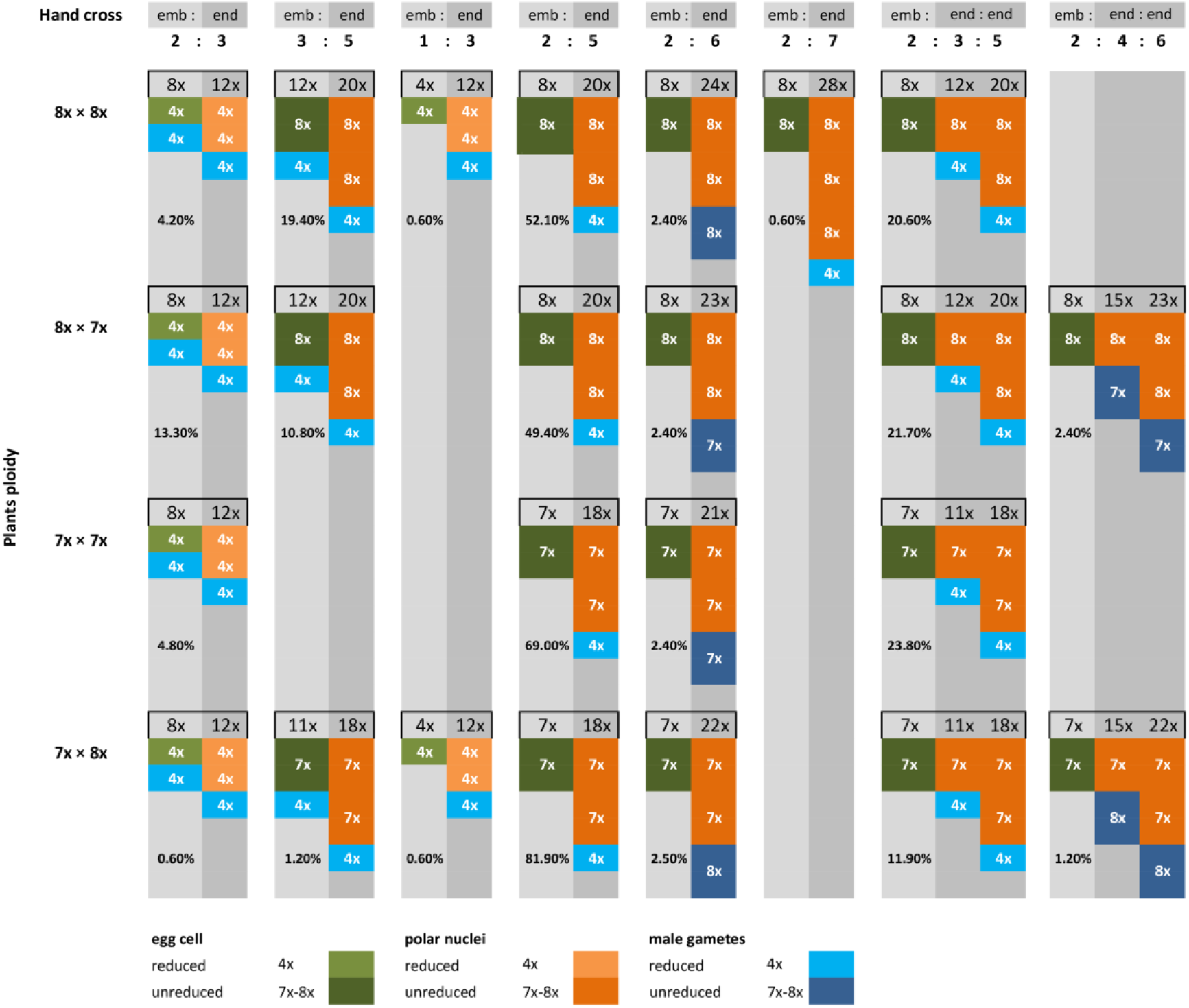
Embryo:endosperm ploidy balances (or embryo:endosperm:endosperm for cases where nuclei with two different ploidy levels coexist in the endosperm), in *Psidium cattleyanum* seeds obtained from hand-pollinated crosses between heptaploid (7x) and octoploid (8x) plants. In each cross, the reduced (4x) or unreduced (7x and 8x) condition of the embryo sac and male gametes and the presence of two or three polar nuclei are considered. emb: embryo, end: endosperm.

The maternal-to-paternal (m:p) contribution to endosperm formation was diverse. In addition to the expected 2m:1p ratio (reduced polar nuclei and male gamete), ratios of 4m:2p (unreduced polar nuclei and male gamete) and 4m:1p (unreduced polar nuclei and reduced male gamete) were also recorded. The 2m:1p contribution occurred in only 5.1% of cases, 4m:2p was present in only 2.4%, and the most frequent was the 4m:1p ratio in 73.0% of cases. To this scenario of diverse contributions, variability was added by the presence of three polar nuclei in unreduced female gametophytes in 19.5% of the analyzed cases. One record showed a 6m:1p contribution when the three unreduced polar nuclei were fertilized with a reduced male gamete (0.2%). In other cases, endosperms were formed by cells with different m:p contributions, such as (4m:1p) + (2m:1p) or (4m:2p) + (2m:2p) when two polar nuclei were fertilized with one of the male gametes and the third with the other male gamete, with male gametes being reduced (18.5%) or unreduced (0.8%), respectively. The diversity in ploidy balances between embryo and endosperm is further influenced by the ploidy levels of the plants, the presence of reduced and unreduced gametes, and the presence of two or three polar nuclei involved in the crosses as shown in Figure 4.

## DISCUSSION

Flow cytometric analysis of single *Psidium cattleyanum* seeds, obtained from controlled crossings, allowed us to determine the embryo:endosperm balance, interpret the origin of each of these two structures, and infer the reproductive pathway by which each seed was formed. The primary reproductive mode in *P. cattleyanum* is facultative pseudogamy with an embryo:endosperm balance of 2:5, but it also exhibits a wide variety of other possible combinations. This diversity is attributed to the presence or absence of double fertilization between pollen grains with reduced or unreduced male gametes, combined with reduced or unreduced embryo sacs that may contain two or three polar nuclei in the central cell. In addition the presence of both even and odd ploidy levels in the studied plants resulted in different ploidy levels in the reduced gametes. All of these factors together explain the high diversity of embryo:endosperm DNA content balances found in this study.

### Diversity in the pathways of embryo and endosperm formation

The 2:5 balance between embryo:endosperm ploidy levels was the most frequent in the analyzed seeds of *Psidium cattleyanum* (64.2%) and corresponds to seeds of pseudogamous apomictic origin (Matzk *et al*., 2000), which explains the morphological and molecular uniformity previously found in studies of seed-derived offspring (Ellshoff *et al*., 1995; Vignale & Bisio, 2005). The 2:5 balance is generally the most reported combination in pseudogamous apomictic species. In this case a pollination stimulus is necessary, and a triple fusion occurs more or less regularly between a reduced male gamete and two unreduced polar nuclei, while the embryo is formed by diploid parthenogenesis (Maheshwari, 1950; Bicknell & Koltunow, 2004). However, we observed that this balance together with a variety of other combinations increases the frequency of apomixis in *P. cattleyanum* to up to 86.5%. In 19.5% of the pseudogamous cases, unreduced embryo sacs with three polar nuclei were present. After fertilization of the central cell, this resulted in an endosperm composed of cells with different ploidy levels (balances 2:5:3 and 2:6:4). The occurrence of different numbers of polar nuclei has already been reported in embryo sacs of other apomictic species (see Nogler, 1984 for examples). Moreover, three polar nuclei were identified in ontogenetic studies of the female gametophyte of *P. cattleyanum* by Souza-Pérez and Speroni (2017). Almost all cases of apomixis in our study corresponded to recurrent apomixis events, where the embryo is formed from an unreduced female gametophyte, usually through diploid parthenogenesis (Maheshwari, 1950; Bicknell & Koltunow, 2004). This process typically results in stable and viable embryos (Raghavan, 1997). However, also a low number of haploid embryos was detected (0.4%), originating from haploid parthenogenesis from a reduced egg cell. These cases of non-recurrent apomixis are rare and produce unstable embryos, which in most cases result in sterile plants (Maheshwari, 1950). However, considering the polyploid status of *P. cattleyanum*, this form of apomixis (polyhaploid parthenogenesis) may be one of the ways to generate changes in plant ploidy levels, as we will discuss later.

Despite the fact that the vast majority of seeds are of apomictic origin, sexuality was detected in 13.5% of the seeds. Generally, most apomictic species are facultative and retain some degree of sexuality (Raghavan, 1997), as observed in various studies (Bicknell & Koltunow, 2004; Cornaro *et al*., 2023; Hörandl, 2022; Reutemann *et al*., 2022). In *P. cattleyanum*, sexual events were detected in both even and odd ploidy mother plants. However, significant differences (p=0.0158) were found in the frequency of sexual events between even-ploidy (24%) and odd-ploidy (3%) mother plants, with 8x plants showing eight times more sexual events than 7x plants. The prevalence of apomixis among odd cytotypes would signify an escape from sterility, as observed in other plant species (Sailer *et al*., 2020). In the case of *P. cattleyanum*, genetic diversity studies using SSR markers in wild populations in Brazil showed higher diversity in populations with even ploidy plants (6x and 8x) than in those with odd ploidy plants (7x) (Machado *et al*., 2020). These authors suggested that the results are linked to the coexistence of apomixis and sexuality in wild populations. In our study, the quantification of both reproductive pathways confirms this hypothesis and provides an accurate and efficient method to explain the origin of diversity within or between populations or agamic complexes.

In facultative apomictic species, both apomixis and sexuality can occur in the same plant (Nogler, 1984; Aliyu *et al*., 2010; Dobeš *et al*., 2013). In our study, we were able to verify that both reproductive pathways can coexist in seeds from the same fruit. Similar results were observed in progenies obtained from seeds from natural populations in Brazil, where different ploidy levels ranging from triploid to decaploid and dodecaploid were obtained from the same fruit (Machado & Forni-Martins, 2022). While the control of apomixis remains unclear, there is recognition of the interplay between genetic, epigenetic, and environmental factors (Terzaroli *et al*., 2023). It has been postulated that temperature fluctuations and light availability can affect reproductive behavior (Asker & Jerling, 1992) and serve as natural triggers for meiotic expression (Hojsgaard & Hörandl, 2019). However, our data confirm that both reproductive pathways are occurring in the same flower, under the same environmental conditions in which flowering occurs, suggesting that in *P. cattleyanum*, this may not be a determining factor in the control of apomixis, and it may instead be related to epigenetic effects.

Of the instances of sexuality events found in *Psidium cattleyanum* (13.5%), approximately one-third (4.7%) correspond to fertilizations of reduced embryo sacs with reduced male gametes. These percentages align with the low frequency of meiosis observed in the megaspore mother cell ontogeny of the female gametophyte in this species (Souza-Pérez & Speroni, 2017). Strikingly, the remaining two-thirds (8.8%) correspond to the fertilization of unreduced egg cell in unreduced embryo sacs, which, like haploid parthenogenesis, are ways of introducing changes in the ploidy levels of the offspring. *P. cattleyanum* embryos with chromosomal content higher than their parents (balance 3:5) were formed by fertilization of an unreduced egg cell. This is consistent with observations in natural populations of other species (Schinkel *et al*., 2017; Macková *et al*., 2020), where it is more common for an unreduced female gamete to be fertilized by a reduced male gamete than the reverse (Harlan & De Wet, 1975; Krahulcová *et al*., 2004). The 3:5 balance, corresponding to this type of sexual event (Matzk *et al*., 2000), occurred at higher frequencies in octoploid mother plants (19.4% and 10.8%) than in heptaploids (1.2%). In crosses performed in hexaploid species of *Hieracium*, this balance was found at a frequency of 3.98% by Krahulcová *et al*. (2004). These variations may be due to the fitness of the mother plants to generate unreduced gametes and their ability to accept fertilization. The occurrence of unreduced male gametes was observed in a low proportion (3.2%), compared to other studies in *P. cattleyanum* where, based on cytological analyses, an average of 8.5% was found for three triploid cytotypes and 31.3% for a tetraploid cytotype (Barbosa, 2016). Although the formation of unreduced gametes has been documented in several genera (Harlan & De Wet, 1975; Bretagnolle & Thomson, 1995; Kreiner *et al*., 2017), most studies have focused on cultivated species. The few studies conducted on wild species have also shown low production of 2n gametes, ranging from 0.1-2%, although some may exceed 10%, (Ramsey 2007; Kreiner *et al*., 2017). It has been reported that the frequency of unreduced gametes varies among species and cultivars (Kreiner *et al*., 2017), and even between flowers of the same plant and within the locules of the same anther (Veilleux *et al*., 1982), similar to what our studies suggest in *P. cattleyanum*, where variability in the generation of this type of gametes within the same flower was observed.

The male gametes involved in fertilization, both of the egg cell and the polar nuclei, were mostly reduced (96.8%). The reduction in the case of pollen donor plants with even ploidy levels (8x) resulted in gametes with a DNA content of 4x, while surprisingly, in pollen donor plants with odd ploidy levels (7x), the DNA content in those gametes was also similar to 4x. Meiotic studies of microspore mother cells, which could provide information on the causes of these frequencies, are rare in the genus *Psidium*, possibly due to the small size of its chromosomes (1.41 to 2.7 µm) and their high number (Vázquez, 2014). In a heptaploid (7x) accession of *P. coriaceum* (syn. *P. cattleyanum*), a high frequency of abnormal meiosis was observed, with the majority forming univalents (20.6II+35.7I), and 80% pollen sterility (Singhal *et al*., 1985). In the case of an octoploid (8x) plant of the same species, mostly bivalents were formed in metaphase I, although univalents and multivalents were also observed (Singh, 2005). Studies on microsporogenesis and pollen grain formation carried out by Souza-Pérez *et al*. (2021), on the same 7x and 8x material used in our study, showed a high number of abnormalities during late meiosis related to a reduction in the number of microspores and heterogeneity in shape, size, and viability of future pollen grains. These abnormalities were detected in both the 8x and 7x materials, but more frequently in the odd ploidy plants, as reflected in the viability values reported (Souza-Pérez *et al*., 2021). Thus, the DNA content similar to 4x found in reduced male gametes of the analyzed 7x materials shows a trend towards the selection of more euploid and balanced pollen grains to overcome the irregularities in odd-polyploid meiosis.

### Maternal and paternal contributions to endosperm formation

The formation of the endosperm in all cases resulted from the fertilization of polar nuclei (pseudogamy). Cases were identified in which both reduced and unreduced gametes and polar nuclei were involved, as well as a combination of both. This led to the formation of endosperms composed of cells with different degrees of ploidy, and more importantly, a wide diversity of embryo:endosperm balances. In total, seven and eight different types of balances between embryo and endosperm were recorded in *Psidium cattleyanum* for heptaploid and octoploid mother plants, respectively, demonstrating a remarkable plasticity in the species. Although uncommon, similar records exist in other species, such as the herbaceous *Hypericum perforatum*, which has 11 identified balances (Matzk *et al*., 2001).

In pseudogamy, double fertilization is excluded, and only single fertilization of the central cell occurs to stimulate endosperm development (Rutishauser, 1982). Therefore, if the second male gamete is not used to fertilize the egg cell, it may be lost or may also contribute to endosperm formation (Talent & Dickinson, 2007). In *Psidium cattleyanum,* our results revealed that in the cases where two fertilizations of polar nuclei occurred, fertilization of the egg cell did not take place. The alteration in the fate of the second male gamete appears to represent a mechanism to circumvent seed incompatibility, a phenomenon observed in other plants with pseudogamous gametophytic apomixis (Nogler, 1984; Savidan, 2000). In the genus *Amelanchier*, a fusion of reduced polar nuclei occurs before pollination in sexual species, similar to the typical double fertilization of Angiosperms. However, in apomictic species of the genus, no fusion was observed, facilitating the independent fertilization of the two polar nuclei with the two male gametes (Campbell *et al*., 1987; Burgess *et al*., 2014). In our single seed analysis, we found in 19.3% a formation of the endosperm by nuclei with different ploidy levels (balances 2:3:5 and 2:4:6), indicating the presence of a third polar nucleus not fused with the other two polar nuclei. This third nucleus is fertilized by the second male gamete, which does not participate in the egg cell fertilization. In some species, the number of polar nuclei and male gametes can vary. Therefore, the ploidy level of the endosperm in pseudogamous species is much more variable than in sexual species (Rutishauser, 1982). The presence of three polar nuclei in the embryo sac has been observed in various plant species (Rutishauser, 1982; Talent & Dickinson, 2007; Lepší *et al*., 2019). In *Ranunculus auricomus* (Rutishauser, 1982), up to three polar nuclei were also identified, and it was observed that they could be fused or not. In addition, it was noted that they could be fertilized by one or two male gametes, generating endosperms with cells of more than one ploidy level in some cases, similar to the flow cytometric results observed for *P. cattleyanum*. However, it is unknown whether the seeds of all the cytological variations found in *R. auricomus* (Rutishauser, 1982) and *P. cattleyanum* germinate successfully or not.

In the shrub species *Sorbus*, the number of polar nuclei varies according to the ploidy of the individuals analyzed, with a proportion of 1% in diploid seeds, 2% in triploids, 3% in tetraploids, and 5% in pentaploids (Lepší *et al*., 2019). In *Psidium cattleyanum*, previous anatomical studies on the same plants we analyzed reported the presence of three polar nuclei in some embryo sacs (Souza-Pérez & Speroni, 2017). Our work reveals that the proportion of seeds with this condition was significantly higher than that recorded in *Sorbus*, with 16.8% of seeds from heptaploid mothers and 22.2% of seeds from octoploid mothers. However, it follows the same trend detected in species of that genus, with a higher frequency of three polar nuclei as the ploidy level increases.

The success in embryo development in most species is largely attributed to a normal development of the endosperm (Brink & Cooper, 1947). Most angiosperms require an accurate balance between maternal and paternal genome contribution of 2:1 to normally develop. Any deviation from this ratio normally leads to seed abortion (Johnston *et al*., 1980; Nogler, 1984). An excess of either maternal or paternal dosage can lead to a failure of seed viability (Kolher *et al*., 2010; Hörandl, 2022). However, the evolution of apomixis involves changes in DNA contribution, leading to alterations in accepted dosages (Nogler, 1984). It has been observed that a 2:1 ratio is more easily modifiable in interploid crosses between low-ploidy plants than between high-ploidy plants, although little is known about the role of imbalance in endosperm development in higher ploidies (Hörandl, 2022). In apomicts where the maternal contribution to the endosperm is 2x or 4x, the 2:1 ratio seems to be strictly necessary. However, once the maternal contribution exceeds the 4x value there is no longer a requirement for endosperm formation (Quarín, 1999). In *Psidium cattleyanum*, flow cytometry revealed different dosages: 2m:1p (5.1%), 4m:1p (73.0%), and 4m:2p (2.4%). Each of these ratios is associated with different events: the first is the expected ratio for cases of sexuality and non-recurrent displospory (both cases with a reduced embryo sac); the second occurs in cases of pseudogamous diplospory and sexuality with a unreduced embryo sac and reduced male gamete; and the last, a case of pseudogamous diplospory with an unreduced male gamete. Nevertheless, there are reports of this last type of dosage in *Arabis holboelli* (Nogler, 1984), maintaining the 2m:1p ratio (Johnston *et al*., 1980). In *Tripsacum dactyloides* and *T. maizar*, functional endosperms are also formed with various contributions from both mother and father, including 2m:1p, 4m:1p, or 4m:2p in sexual plants and 8m:1p or 8m:2p in apomicts (Grimanelli *et al*., 1997). *Tripsacum* species, like *P. cattleyanum*, do not show specific dosage requirements for endosperm formation, suggesting that the evolution of diplospory apomixis might be restricted to species with few or no dosage requirements (Grimanelli *et al*., 1997). However, it is interesting to note that in our study, the highest frequencies recorded correspond to ratios of 4m:1p and 2m:1p, and combinations of both in cases of the presence of three polar nuclei and independent fertilizations to form a mosaic of cells with these ratios.

### Relevance of our findings for wild populations

Ninety percent of the genus *Psidium* is composed of species or cytotypes with a polyploid origin. Although odd ploidy levels are rarer, they are often associated with individuals exhibiting irregularities in sexual reproduction due to meiotic irregularities (Costa, 2009). *Psidium cattleyanum* displays a broad range of cytotypes in wild populations (Atchinson, 1947; Singhal *et al*., 1985; Raseira & Raseira, 1996; Costa & Forni-Martins, 2006; Vázquez, 2014; Machado *et al*., 2020; Machado & Forni-Martins, 2022). Embryos from seeds analyzed in our study, formed by apomictic or sexual reproduction, showed ploidy levels of tetraploids, heptaploids, octoploids, undecaploids, and dodecaploids. The undecaploid (2n=11x=121) represents a novel observation for this species.

Our studies revealed the formation of unreduced female gametes and reduced male gametes at a high frequency. The formation of unreduced female gametes and sexual hybridization (2n+n) represent the major route for inducing polyploids, and explains the presence of different cytotypes in the same population. In our study, the high ploidy levels (11x and 12x) found in the analyzed embryos align with this frequent route of formation from unreduced embryo sacs (2C+C) fertilized by reduced male gametes. The detected unreduced male gametes occurred at a low frequency (3.2%) and were associated with apomictic events in 2:6 and 2:4:6 balances, originating from a parthenogenetic embryo development, and thus did not contribute to increased embryo ploidy levels. The low proportion of unreduced male gametes compared with female gametes aligns with results obtained for other tree genera such as *Sorbus* (Lepší *et al*., 2019) and support the general premise that apomixis primarily involves unreduced female gametes, whereas microsporogenesis meiosis remains unaffected, favoring the subsequent production of reduced male gametophytes (Asker & Jerling, 1992; Lepší *et al*., 2019).

In the analyzed seeds, we found a high frequency (86.1%) of diploid parthenogenesis and a 0.4% occurrence of tetrahaploids (4x) resulting from polyhaploid parthenogenesis (haploid parthenogenesis of a polyploid). Therefore, the embryo:endosperm DNA content balance is 1:3 (Matzk *et al*., 2000). The process of polyhaploid parthenogenesis involves meiosis (recombination exists) but avoids fertilization. The embryo is formed by parthenogenesis and has half the chromosomal load of the mother plant. Although polyhaploid parthenogenesis is observed in low proportions in natural populations and is generally not very successful (Asker & Jerling, 1992), reciprocal crosses between *Pilosella rubra* (apomictic 6x) and *P. officinarum* (sexual 4x) resulted in a high frequency of polyhaploid progeny (Rosenbaumová *et al*., 2012). The formation of polyhaploids could explain the origin of some odd ploidies in *Psidium cattleyanum*, such as cases where triploid and hexaploid or tetraploid and octoploid offspring coexist in seeds from the same fruit collected from wild populations in Brazil (Machado & Forni-Martins, 2022). It is necessary to confirm through population studies whether these cases of polyhaploid seedlings or embryos and the wide range of ploidy values detected at this level persist in nature. This would enhance our understanding of the occurrence and nature of polyploids and their contribution to the evolution of angiosperms (Heslop-Harrison *et al*., 2023). The report of polyhaploids in *P. cattleyanum* also confirms that, although tetrads of megaspores were not observed in ontogenetic studies by Souza-Pérez & Speroni (2017), meiosis occurs at a low frequency. The majority of female gametes were unreduced (94.9%), supporting observations of low meiosis occurrence and that *P. cattleyanum* is predominantly an apomictic species.

## CONCLUSIONS

The different pathways of seed formation identified in this study indicate that *Psidium cattleyanum* is predominantly an apomictic species (approximately 87%), maintaining a margin of sexuality (facultative apomixis). Almost all cases of detected apomixis are of the recurrent type (approximately 99.5%) with embryos showing high potential viability. The sexual pathway with reduced embryo sacs was more common in plants with even ploidy than in those with odd ploidy level. Only about a third of sexual reproduction cases occurred in reduced embryo sacs. Fertilization of unreduced embryo sacs, along with cases of non-recurrent apomixis (although in low proportion), represents two pathways for generating ploidy changes in polyploid species. In this pseudogamous species, the endosperm is formed from different maternal-paternal contributions, and the coexistence of cells with different ploidy levels is observed. While these observations confirm the plasticity of this ephemeral tissue, there is a greater tendency toward doses of 2m:1p (5.1%), 4m:1p (73%), and their combinations (18.5%) in the cells of the endosperm. Our results may be the basis for exploring the frequency of apomixis and sexuality in wild populations and the persistence of the detected ploidy changes in nature.

## COMPETING INTERESTS

There is no conflict of interest to declare by the authors.

## AUTHOR CONTRIBUTIONS

Da Luz-Graña, C. contributed to conceptualization, flow cytometry analysis, original draft writing, editing and visualization. Vaio, M. contributed to conceptualization, guided the flow cytometry analysis, original draft writing, editing and visualization. Fuchs, J. guided C.D.LG in optimization of the flow cytometry technique and contributed to data interpretation, reviewing, and manuscript editing. Borges, A. performed the statistical analysis, contributed in the review, and manuscript editing. Speroni, G. contributed to conceptualization, writing the original draft, editing, visualization, project administration, and acquisition of funding. All authors read and approved the final manuscript.

## ACKNOWLEDGEMENTS

We thank Comisión Sectorial de Investigación Científica (CSIC), I+D projects, Universidad de la República, Uruguay, for granting the financial support. C.D.LG. thanks Postgraduate program-Facultad de Agronomía in which she did her Master’s Degree studies, Comisión Académica de Posgrado-Universidad de la República by grants scholarships and the Leibniz Institute of Plant Genetics and Crop Plant Research where she conducted the laboratory training for optimizing the flow cytometry technique in seeds.

## DATA AVAILABILITY

The data underlying this article will be shared on reasonable request to the corresponding author.

## Notes

### Competing Interest Statement

The authors have declared no competing interest.

